# Inhibition of SHP-1 /2 blocks antigen cross-presentation by human macrophages and dendritic cells

**DOI:** 10.1101/2025.10.21.683666

**Authors:** Alexine S. de Wit, Harry Warner, Huang Huang, Martijn Verdoes, Geert van den Bogaart, Frans Bianchi

**Affiliations:** Department of Molecular Immunology, Groningen Biomolecular Sciences and Biotechnology Institute, University of Groningen, Groningen, the Netherlands; Department of Infectious Diseases, Huashan Hospital, Institute of Infection and Health, Fudan University, Shanghai, China; Department of Tumor Immunology, Radboud Institute for Molecular Life Sciences, Radboud University Medical Center, 6525 GA Nijmegen, the Netherlands; Institute for Chemical Immunology, Radboud Institute for Molecular Life Sciences, Radboud University Medical Center, 6525 GA Nijmegen, the Netherlands

## Abstract

PD-1 immune checkpoint therapy aims to stimulate T-cell responses against cancer, but faces challenges due to resistance, rendering it ineffective for a significant subset of patients. Inhibitors of SHP-1 and SHP-2, widely expressed protein tyrosine phosphatases known for their pro-cancer and immunosuppressive properties, have attracted attention for their potential to enhance therapy efficacy and overcome resistance when combined with immune checkpoint PD-1 blockade. However, how SHP-1/2 inhibition affects antigen presenting cells is incompletely understood. In this study, we evaluated the effect of SHP-1/2 inhibition on antigen cross-presentation by human monocyte-derived macrophages and dendritic cells, using T cell reporter cell lines specific for epitopes derived from cancer antigens NY-ESO-1 and gp100. Our findings indicate that SHP-1/2 inhibitor NSC-87877 significantly reduces the cross-presentation efficiency of both antigens. Mechanistically, we show that SHP-1/2 inhibition blocks endo/lysosomal acidification and the activation of cathepsin proteases. The reduction of antigen cross-presentation upon SHP-1/2 inhibition potentially limits the effectiveness of the combination therapy with immune checkpoint inhibition.

## Introduction

Cytolytic CD8^+^ T cells (CTLs) play key roles in the immune responses against cancer, viruses and other intracellular microbial pathogens. CTLs recognize malignantly transformed or infected host cells by a process called antigen presentation. Here, the host cells present peptide fragments derived by proteolysis from proteins selectively expressed by pathogenic microbes or cancer cells, so-called epitopes, on their major histocompatibility complex (MHC) class I to the effector CTLs. This promotes the killing of the infected or malignant cells by the CTLs. However, in order to become effector CTLs, naive CD8^+^ T cells need to be activated by dendritic cells (DCs) in the secondary lymphoid organs [1]. This process occurs through antigen cross-presentation, a process where DCs present exogenous antigens on their MHC class I. In addition, macrophages and other immune cells are also capable of antigen cross-presentation, and this likely functions for the re-activation of effector and memory CD8^+^ T cells [2].

In T cells, the binding of the T-cell receptor (TCR) to a cognate MHC-epitope complex triggers the phosphorylation of signaling proteins by multiple kinases. At later stages of TCR signaling, these include serine/threonine kinases (e.g. protein kinase C) and phosphoinositide 3-kinases, but TCR signaling is initiated by tyrosine kinases. First, the tyrosine kinase LCK phosphorylates immunoreceptor tyrosine-based activation motifs (ITAM) on the CD3 subunits of the TCR complex and downstream adapters. ITAM phosphorylation in turn results in recruitment and activation of other tyrosine kinases (e.g. ITK, ZAP-70). However, this tyrosine phosphorylation is countered by inhibitory signaling by programmed cell death protein 1 (PD-1). PD-1 is a membrane protein that binds to its receptors PD-1 ligands 1 and 2 (PD-L1 and PD-L2) on the surface of antigen presenting cells. PD-1 contains an immunoreceptor tyrosine-based inhibition motif (ITIM) motif that can recruit Src homology region 2 domain-containing phosphatase-2 (SHP-2), also known as tyrosine-protein phosphatase non-receptor type 11 (PTPN11), and potentially also SHP-1 (known as PTPN6) [3], through their SH2-motifs [4, 5]. SHP-1/2 are tyrosine phosphatases that directly counter T cell receptor signaling and thereby prevent T cell activation [6]. For example, studies with primary human T cells showed that SHP-2 inhibits T cell function through dephosphorylation of the kinase ITK [7].

PD-1 and its ligands are targets of so-called immune checkpoint inhibitor (ICI) therapy [8]. This antibody-based therapy is in the clinic for various cancers, including melanoma and non-small cell lung cancer, renal cell carcinoma, bladder cancer, Hodgkin’s lymphoma, and head and neck squamous cell carcinoma. Administration of PD-1 or PD-L1 binding antibody in cancer patients prevents PD-1 binding to PD-L1 through steric hindrance and/or competitive binding, thereby promotes T-cell activation. This enables the immune system to recognize and attack cancerous cells more effectively. PD-1 ICI therapy has shown remarkable success in treating the above-mentioned cancers, often offering durable responses. However, the effectiveness of checkpoint immunotherapy varies widely among patients, and response rates to PD-1 ICI therapy can be as low as 20% for certain cancer types [8]. Therefore, there is an ongoing search for combination therapies that can increase the efficiency of PD-1 ICI therapies.

One of the suggested combination therapies is PD-1/PD-L1 ICI therapy in combination with inhibitors for SHP-1 and/or SHP-2. Both phosphatases are well recognized therapeutic targets in cancer, and their heightened expression and/or abnormal activation have been reported for diverse solid tumors and hematological malignancies [9, 10]. Especially SHP-2 plays critical roles in regulating various cellular processes downstream of tyrosine kinase signaling pathways, including cell growth, survival, differentiation, and migration [10]. Therefore, small molecule inhibitors targeting SHP-2 are being developed for cancer treatment. These inhibitors disrupt SHP-2 activity, thereby impeding the aberrant signaling pathways that propel cancer progression. Clinical trials are presently underway to assess the safety and efficacy of SHP-2 inhibitors either alone or in combination with other cancer therapies across various cancer types [11-13].

The combination of SHP-1/2 inhibitors with PD-1 ICI therapy could be very potent, because SHP-2 counters T-cell activation and can make ICI therapy more effective [14, 15]. Moreover, SHP-2 promotes a more immunosuppressive tumor microenvironment by increasing the production of immunosuppressive cytokines and chemokines, and by preventing the differentiation of infiltrating myeloid cells into effector cells [16]. SHP-2 also promotes cancer progression via the activation of the RAS/RAF/ERK/MAPK pathway [17]. Because of these reasons, SHP-2 is implicated in the resistance to ICI therapy [11]. Indeed, preclinical studies showed that inhibiting SHP-2 can increase the efficacy of ICI therapy by reducing the immunosuppressive tumor microenvironment, increasing antigen presentation by cancer cells, and promoting the clearance of cancer cells [14, 18, 19]. Moreover, the combination of SHP-2 inhibitors with PD-1 ICI therapy is more effective in inhibiting tumor development than the single treatment alone [18]. There are ongoing phase I and II clinical trials testing SHP-2 inhibitors together with PD1 or PD-L1 ICI therapy [11, 12].

Interestingly, in our recent phosphoproteomics study of human monocyte-derived DCs, we found that genes involved in antigen presentation undergo phosphorylation or dephosphorylation following stimulation with the canonical immune stimulus lipopolysaccharide (LPS) [20]. Analyzing samples from six different donors, we identified 2,068 phosphosites that showed significant changes in phosphorylation levels at 1 hour and/or 4 hours after LPS exposure. These included proteins such as hydrolases, the transporter associated with antigen presentation (TAP), the V-ATPase, MHC class I, and the NADPH oxidase NOX2. Of the 2,068 altered phosphorylation sites, the vast majority were phosphorylated threonines and serines, and only 12 (i.e., 0.5%) were phosphorylated tyrosines (Figure 1). Six of these significantly altered phosphotyrosines were upregulated at 1 hour after pathogenic stimulation, indicating that they became phosphorylated by unknown tyrosine kinases. It might well be that SHP-1/2 inhibition further increases their phosphorylation, potentially affecting cross-presentation. Although the functional consequences of most of these phosphorylation and dephosphorylation events remain unclear, SHP-1/2 inhibitors may modulate them, thereby affecting cross presentation efficacy and as such impacting effective CTL populations. This could potentially impair the efficacy of ICI therapy. In this study, we evaluated the effect of SHP-1/2 inhibition on antigen cross-presentation by human monocyte-derived macrophages and DCs. Using T-cell reporter cell lines specific for epitopes derived from two well-known cancer antigens, we demonstrated a significantly reduced cross-presentation efficiency of both antigens.

**Figure 1.**
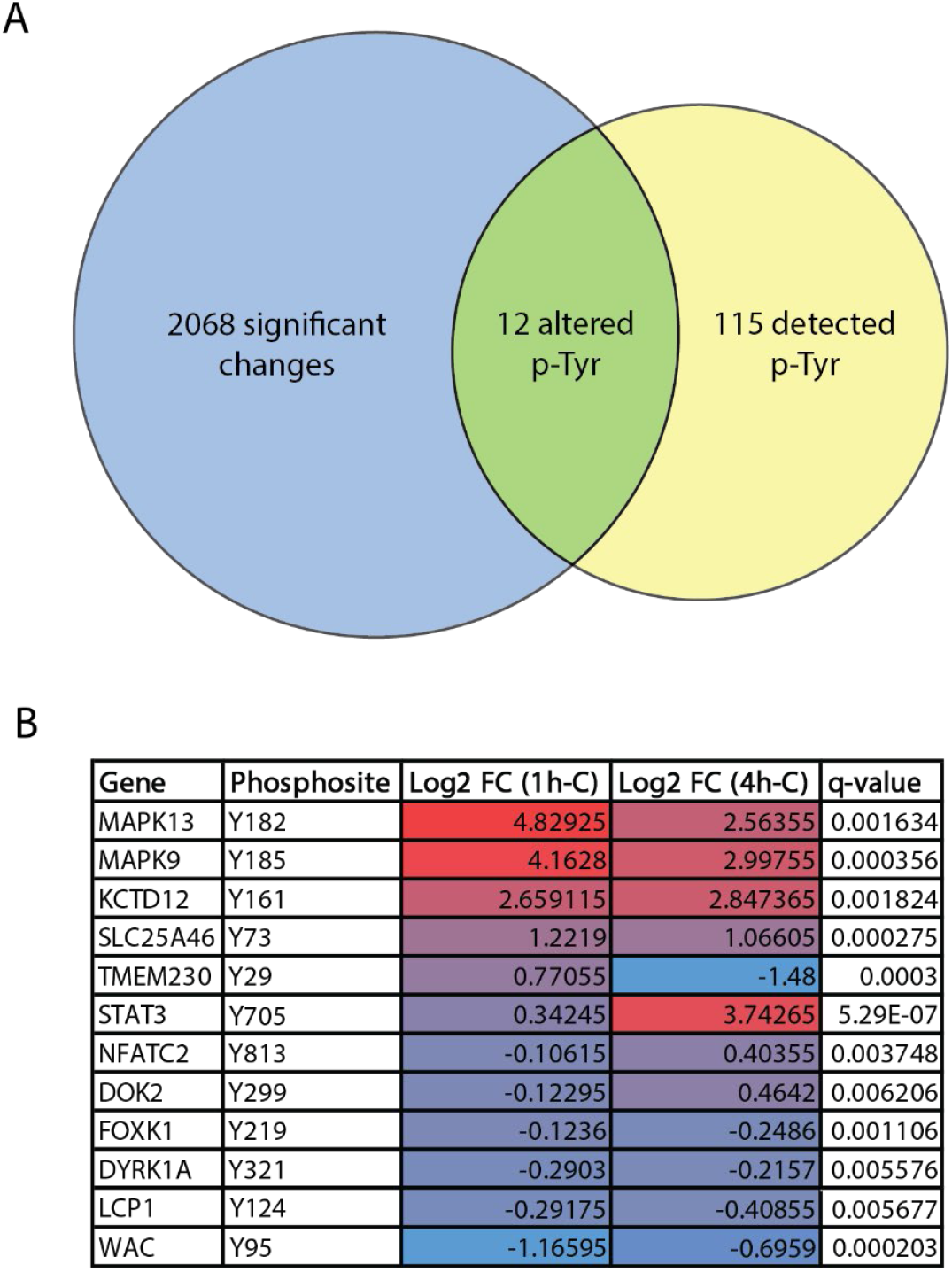
Activation of dendritic cells triggers protein phosphorylation and dephosphorylation. Analysis of our previously published phosphoproteomics data from human monocyte-derived DCs from 6 different donors that were stimulated for 1 or 4 hours with LPS [25]. (A) Venn diagram showing the numbers of significantly altered phosphorylation sites and of detected phosphorylated tyrosines. (B) Table listing the 12 significantly altered phosphorylated tryosines. FC: fold change; p-Tyr: phospho-tyrosine

## Materials and methods

### Antigen presenting cells

Buffy coats of healthy HLA-A2^+^ donors were obtained from the Dutch blood bank. The utilization of human blood received approval from the Dutch blood bank, with adherence to national and institutional guidelines during all experiments. Informed consent was obtained from all donors by the blood bank. Samples were anonymized, and the investigators could not determine the identity of the blood donors.

Peripheral blood mononuclear cells (PBMC) were isolated from buffy coats using Lymphoprep (Stemcell Technologies; 07861)-mediated density centrifugation followed by magnetic bead isolation of CD14+ (Miltenyi Biotech; 130050201). Monocyte-derived DCs (moDC) were obtained by culturing CD14^+^ monocytes in complete RPMI-1640 (2 mM L-glutamine (Gibco; 11539876), 10% FBS (HyClone; 12389802), 1% v/v anti-anti (Gibco; 11570486)), supplemented with 300 U/ml interleukin 4 (IL-4; Miltenyi Biotech; 130-093-924) and 450 U/ml granulocyte macrophage colony-stimulated factor (GM-CSF; Miltenyi Biotech; 170-076-136) for 6 days (37 °C and 5% CO_2_). Monocyte-derived macrophages were differentiated by culturing in complete RPMI-1640 supplemented with 100 ng/ml macrophage colony-stimulated factor (M-CSF, R&D Systems; 216-MC) for 7 days (37 °C and 5% CO_2_).

### Jurkat reporter T-cell line

The Jurkat reporter T-cell lines were obtained by lentiviral transduction with plasmids as previously described [21]. To create the Jurkat cells, lentiviral transduction of pNL[NlucP/NFAT-RE/Hygro] (Promega) was performed on Jurkat 76 T-cell line. Gene fragments of TCRα and β chains of TCRs against NY-ESO1 and gp100 coupled with a P2A sequence were cloned into the N103 vector (nLV Dual Promoter EF-1a-MCS-PGK-Puro). Whole length TCR sequences of TCRs against NY-ESO-1_157–165_ and gp100_280-288_ were optimized by implementing known mutations that increase TCR affinity [22-25]. HEK-293T cells were seeded 24 h before transfection in 10-cm dishes at 5*10^6^ cells per dish. Cells were co-transfected with 10 μg of transfer vector, 7.5 μg of envelope vector (pMD2.G), 2.5 μg of packaging vector (psPAX2) and 75 μl PEI (Sigma) to generate lentiviral supernatants. 16 hours post-transduction, culture medium was replaced, and viral supernatants were harvested 48 hours later. The supernatants were filtered (0.45 μm SFCA syringe filter, Corning) and concentrated by centrifugation with a 100 K Amicon Ultra-15 filter (Millipore). Concentrated viruses were used for J76-NFATRE-luc cell transduction using spinoculation for 2 hours in the presence of 6 μg ml−1 polybrene (Sigma). TCR expression was assessed 48 hours after transduction by flow cytometry.

### Antigen presentation assay

To assess antigen presentation, antigen presenting cells (APC) were seeded in a 96-well plate in serum-free RPMI-1640. MoDCs were stimulated with 1 µg/ml lipopolysaccharide (LPS) for 4 hours before the experiment. Cells were pulsed with different concentrations of SHP-1/2 inhibitor NSC-87877 (Sigma-Merck, 565851) and 10 µM of short or long peptide derived from cancer antigens NY-ESO-1 and gp100 (short^NY-ESO-1^: SLLMWITQV; long^NY-ESO-1^: LQQLSLLMWITQVFL; short^gp100^: YLEPGPVTA; long^gp100^: LQQLYLEPGPVTAFL; Genscript). After 3 hours, cells were extensively washed with serum-free RPMI-1640 and Jurkat cells expressing the appropriate TCR were added to the APCs. After 6 hours of coculturing, luciferase activity was measured using the Nano-Glo Luciferase Assay (Promega; N1130) according to the manufacture’s protocol and a plate reader (Synergy HTX, BioTek).

### Assessment of antigen uptake and endosomal acidification

The effect of the inhibitor on uptake and endosomal acidification was assessed by incubating cells (30 min, 37 °C) with 10,000 MW Dextran, Alexa Fluor™ 647 (ThermoFisher; D22914) and 10,000 MW Dextran, pHrodo Green (ThermoFisher: P35368), respectively. The pH calibration curve was prepared as described previously [26]. Cells were treated with with 10,000 MW Dextran, Alexa Fluor 647 and 10,000 MW Dextran, pHrodo Green for 30 minutes at 37 °C. Afterwards, cells were permeabilized and washed in buffers of known pH and analyzed with flow cytometry. Cell viability was determined by incubating cells with eFluor450 (ThermoFisher; 65-0865-18) and cells were analyzed with flow cytometry (Cytoflex, Beckman Coulter).

### Analysis of cathepsin activity

The cathepsin activity was determined by treating cells with NSC-87877 for 1 hour, followed by incubation of the cells with activity-based protease probe BMV109-Cy5 [27] for 30 minutes. Then, cells were lysed in a hypotonic lysis buffer (50 mM PIPES pH 7.4, 10 mM KCl, 5 mM MgCl_2_, 2 mM EDTA, 4 mM DTT, and 1% NP-40) and protein concentration was determined by Pierce BCA protein assay kit (Thermofisher, 23225). Samples were resolved in a 15% SDS-page gel and imaged on an Odyssey XF imager (LiCor). The gel was then blotted and immunostained. The following antibodies were used: rabbit polyclonal anti-Cathepsin S (1:250, Novus Biologicals; NBP2-85807), rabbit polyclonal anti-GAPDH (14C10) at 1:1,000 (1:1000, Cell Signaling; 2118) and goat-anti-Rabbit IgG (H + L) labeled with IRDye 800CW (1: 5000, LiCor; 926-32211). The blot was imaged on an Odyssey XF imager and the image was quantified using ImageJ.

## Results

To address the effects of SHP-1/2 inhibition on antigen cross-presentation on human antigen-presenting cells, we performed cross-presentation assays with NSC-87877 (i.e. the same SHP-1/2 inhibitor as used previously for mouse DCs [24]) and human peripheral blood monocyte-derived macrophages and DCs. We tested the cross-presentation efficiency of two well-studied cancer antigens: the New York esophageal squamous cell carcinoma 1 antigen (NY-ESO-1) and glycoprotein 100 (gp100). The aberrant expression of these proteins in various cancers makes that they can induce anti-cancer immune responses by CTLs. We engineered two Jurkat T-cell lines for stable expression of T-cell receptors that selectively recognize peptide fragments NY-ESO-1_157–165_ and gp100_280-288_ in the context of MHC class I allotype HLA-A*2:01. In addition, the cell lines contained the gene coding for luciferase behind the NFAT promoter, allowing us to detect antigen recognition via a luminescence assay. Importantly, the Jurkat cells only require cognate peptide-MHC complex for activation, which allowed us to discern the antigen cross-presentation efficiency from co-stimulation by CD80/CD86-CD28 interactions and cytokines (i.e., cross-priming).

Before co-culturing with the Jurkat cells, monocyte-derived DCs and macrophages were treated with elongated peptides carrying the NY-ESO-1 or gp100 epitopes. The cross-presentation of these epitopes requires the removal of the extra N- and C-terminal residues by proteolytic cleavage within endosomal or lysosomal compartments or the cytosol. In addition, the DCs and macrophages were treated with the SHP-1/2 inhibitor NSC-87877 for 3 hours. We tested a concentration range from 25 to 200 μM, which did not impact cell viability (Figure 2A). While the inhibitor does not affect luciferase production of the Jurkat reporter T-cell line (Supplemental fig. 1), DCs and macrophages were extensively washed prior to the coculturing with the Jurkat cells.

**Figure 2.**
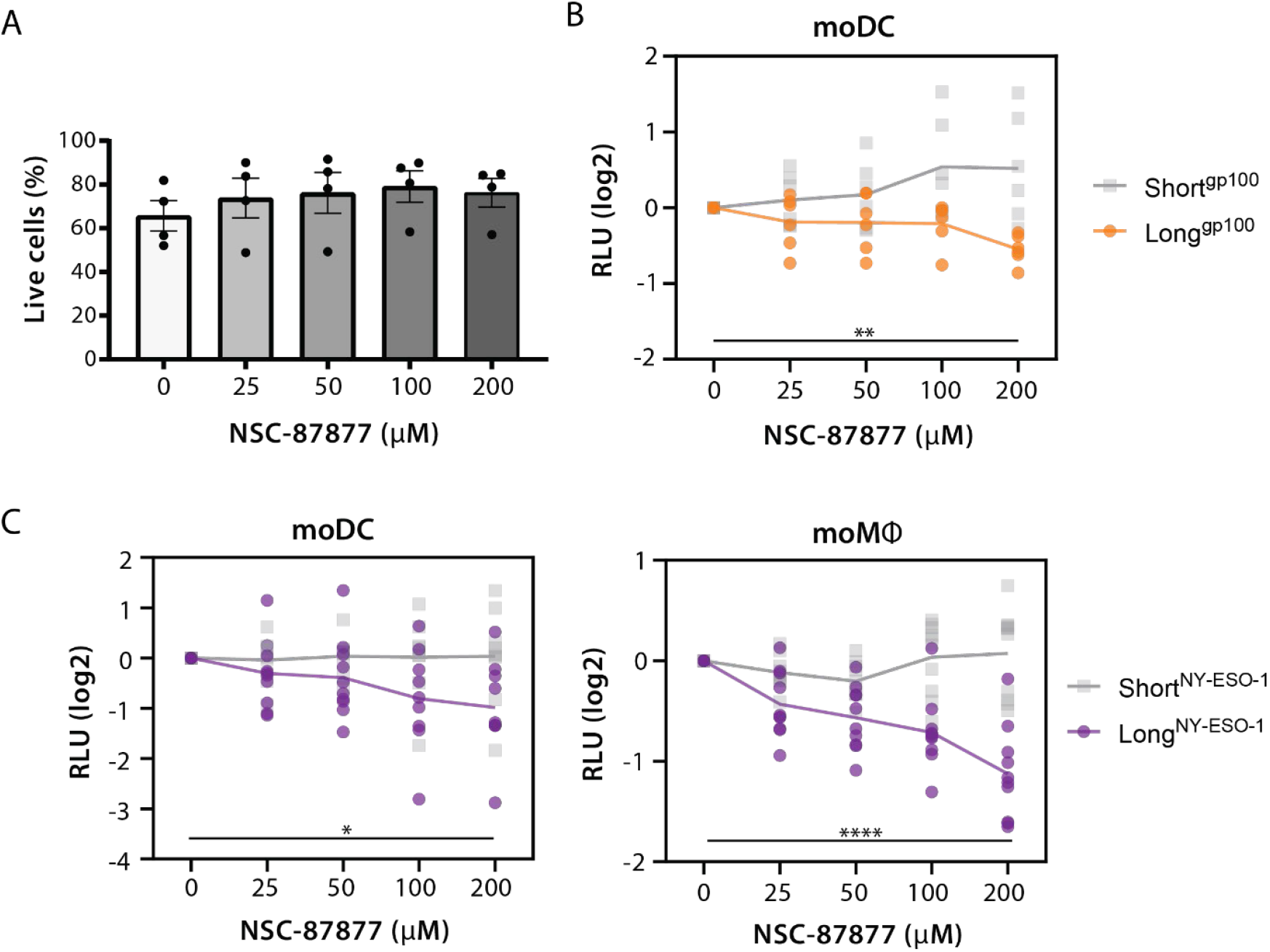
Inhibition of SHP-1/2 leads to reduced antigen cross-presentation. **(A)** Assessment of cell viability by flow cytometry after treatment of monocyte-derived dendritic cells with SHP-1/2 inhibitor NSC-87877 for 3 hours. **(B)** Antigen-presenting cells were treated with different concentrations of NSC-87877 and pulsed with short (short^gp100^) or long (long^gp100^) peptide derived from cancer antigen gp100. Antigen cross-presentation is measured in luminescence (relative luminescence unit, RLU) and normalized against the vehicle (DMSO) condition. **(C)** Same as in panel B, but with short (short^NY-ESO-1^) or long (long^NY-ESO-1^) derived from cancer antigen NY-ESO-1. Statistical significance was determined by performing a one-way ANOVA. A p value < 0.05 was regarded as statistically significant (* p < 0.05, ** p < 0.01, *** p < 0.001, **** p < 0.0001).

SHP-1/2 inhibition lowered cross-presentation efficiency dramatically for both the NY-ESO-1 and gp100 antigens in DCs, and for the NY-ESO-1 antigen in macrophages, in a dose-dependent manner (Figure 2B-C). At the highest concentration, cross-presentation efficiency was decreased up to 2-fold for both cell types compared to the control condition without inhibitor. This reduction was consistently observed for all tested donors. Importantly, NSC-87877 did not reduce the presentation of the short NY-ESO-1 and gp100 peptides, that consist of only the epitope and can be directly presented on MHC class I without further processing in endo/lysosomal compartments or the cytosol.

We also investigated the effect of SHP-1/2 inhibition on uptake of the model antigen dextran by the cells. We observed that uptake of fluorescently labeled dextran by both DCs and macrophages was not affected by treatment with NSC-87877 (Figure 3A). Furthermore, we investigated the effect of SHP-1/2 inhibition on endo-/lysosomal acidification, as SHP-1 has been associated with regulating the maturation and acidification of endosomes [28-30]. We treated cells with pHrodo-labeled dextran, which becomes increasingly more fluorescent with increasing endosomal acidity. After accounting for uptake by treating the cells with dextran labeled with a pH-insensitive fluorophore and normalizing pHrodo intensity accordingly, we generated a pH calibration curve (Supplemental figure 1) to convert fluorescence intensities into pH values. After treatment with NSC-87877, we observed a dose-dependent increase of endosomal pH (Figure 3B), indicating that blockage of SHP-1/2 resulted in inhibition of endo/lysosomal acidification.

**Figure 3.**
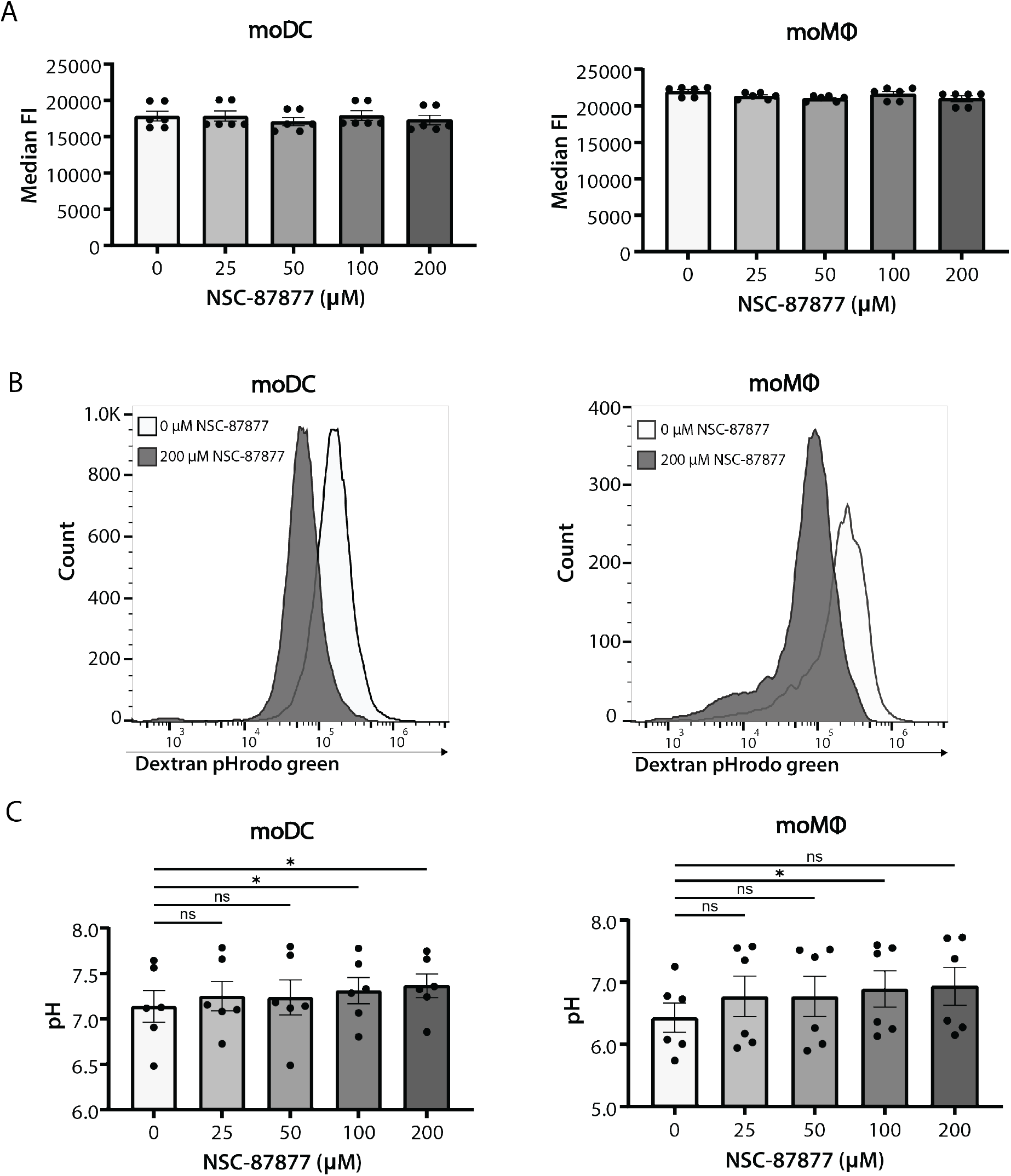
SHP-1/2 inhibitor NSC-87877 does not affect endocytosis but blocks endo/lysosomal acidification. **(A)** Endocytosis by monocyte-derived dendritic cells (moDC, left panel) and macrophages (moMφ, right panel) was determined by treating cells with Alexa Fluor 647-conjugated dextran. **(B)** Representative flow cytometry histograms of moDCs and moMφs treated with pHrodo-conjugated dextran at 0 and 200 µM NSC-87877. **(C)** Endo/lysosomal pH changes in DCs and macrophages after inhibition of SHP-1/2. Cells were treated with pHrodo-conjugated dextran to measure endo/lysosomal pH. Cells were simultaneously treated with Alexa Fluor 647-conjugated dextran to correct for uptake. pH is determined by correcting the pHrodo intensity to uptake and calculating from these intensities by a pH calibration curve. Data from two separate experiments from 6 different donors. Statistical significance was determined by performing a one-way ANOVA. A p value < 0.05 was regarded as statistically significant (* p < 0.05, ** p < 0.01, *** p < 0.001, **** p < 0.0001).

Because endo/lysosomal acidification is a requirement for activation of cathepsin proteases, we also determined the activity and expression of cathepsin S, a phagosomal protease that is associated with antigen cross-presentation [31], of which the activity is dependent on pH. We treated macrophages with 200 µM NSC-87877 (the highest concentration) and BMV109-Cy5, a quenched activity-based protease probe that only becomes fluorescent after cleavage by active cathepsins [32]. This probe covalently binds to the target cathepsin, allowing for detection with in-gel fluorescence [32] (Figure 4A). We found that the inhibition of SHP-1/2 diminished cathepsin S activity by ∼75%, while its expression (determined by western blot) remained unaffected (Figure 4B-C).

**Figure 4.**
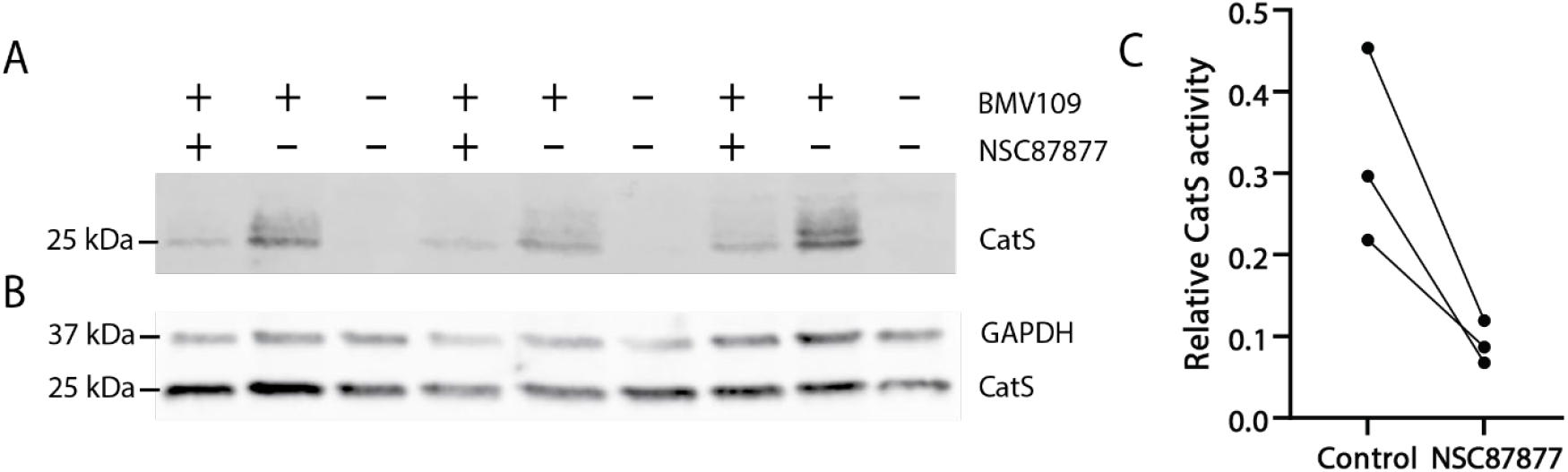
SHP-1/2 inhibitor NSC-87877 blocks activation of Cathepsin S. Analysis of phagosomal protease cathepsin S activity and expression in monocyte-derived macrophages. Cells were treated with or without NSC-87877 and BMV109 activity-based protease probe. **(A)** SDS-page gel with in-gel fluorescence of BMV109 probe. Different heights indicate different cathepsins. **(B)** Western blot showing expression of cathepsin S. GAPDH was used as loading control. **(C)** Quantification of relative activity of cathepsin S. The values are obtained by normalizing cathepsin activity to expression levels. Data from 3 donors individually shown.

## Discussion

In this study, we found that SHP-1/2 inhibition by NSC-87877 blocks antigen cross-presentation. Mechanistically, we showed that SHP-1/2 inhibition results in a reduction of endo-/lysosomal acidification and activation of cathepsin S. How SHP-1/2 affects these processes is not completely understood. It was shown in murine bone marrow-derived macrophages that recruitment of SHP-1 to early endosomes is required for endosomal maturation and acidification [29]. Furthermore, SHP-1 can be recruited to endosomes to induce endosomal acidification by dephosphorylation of NOX2 component p47^phox^, which leads to impaired cross-presentation in murine splenic DCs in a *Listeria monocytogenes*-OVA infection model [30]. In contrast, it has also been shown in human monocyte-derived macrophages that SHP-1 is recruited to phagosomes containing *Mycobacterium tuberculosis* and that this inhibits phagosomal maturation, leading to better bacterial survival [33].

Our observations contrast earlier findings that SHP-1/2 inhibition in mouse CD11c^+^ DCs leads to increased CTL activation [28]. This highlights the complexity of antigen cross-presenting mechanisms and how these could differ between cell types and antigens. The reduction in cross-presentation following SHP-1/2 inhibition might not be a universal phenomenon, because a reduction of endo/lysosomal acidization and cathepsin activation have been reported to also increase antigen cross-presentation. For example, it has been proposed that DCs are good cross-presenters because of their relatively high endosomal pH [34], though others suggest that endosomal acidification is required for cross-presentation of cell-associated pathogens in human DCs [35]. Similarly, while it has been observed that cathepsin S activity is essential for cross-presentation [31], it is also suggested that the decreased activity of cathepsin S, caused by the higher endosomal pH in DCs, is explanatory for the enhanced cross-presentation capacities of these cells [36]. Furthermore, other endosomal proteases of which the activity is also pH-dependent, such as cathepsin D or cathepsin X, and that can contribute to cross-presentation have not been assessed in this study [35, 37]. Consequently, even though we observed the reduction of cross-presentation for two different antigens and for both macrophages and DCs, our findings might not be generally applicable for all antigens and cell types.

Our findings strengthen the notion that SHP-1/2 is important for adaptive immune responses, as has been reported especially in T cells. For example, murine studies with T cell selective knock-in of a constitutively active SHP-2 mutant showed that the numbers of CD8^+^ T cells were reduced in bone marrow and spleen, while the numbers of CD4^+^ cells were increased. Moreover, the numbers of effector memory CD8^+^ T cells were also increased, indicating that SHP-2 promotes T cell memory formation and reduces T cell activation [38]. In addition, SHP-2 not only inhibits but can also promote T cell development and TCR signaling. Conditional knockout of SHP-2 in T cells resulted in a significant block in thymocyte development in the thymus, and cells lacking SHP-2 displayed reduced TCR signaling [39]. Mechanistically, experiments with phosphospecific antibodies showed that SHP-2 promotes TCR signaling through the ERK pathway, and SHP-2 knockout cells showed reduced ERK activation [39]. One possible explanation for how SHP-2 can both promote and inhibit T cell activation might be the association with PD-1. PD-1 recruits SHP-2 to membrane microclusters with the TCR and downstream signaling molecules, where SHP-2 may locally counteract TCR signaling by dephosphorylating activating tyrosines [5]. In contrast, SHP-2 that is not bound to PD-1 might dephosphorylate inhibitory sites on for example ERK regulatory proteins [39], and thereby promote T cell functions.

In addition to its effects on T cells, our findings are in line with previous research showing that SHP-1/2 inhibition also affects other immune cells, which could have implications for PD-1 ICI therapy. For example, SHP-2 has been shown to be important for DC and macrophage responses to fungal infections with the fungal pathogen *Candida albicans* [40]. Knockdown of SHP-2 blocked tyrosine kinase SYK signaling, thereby lowering the production of inflammatory cytokines as SHP-2 recruits SYK to integrins and C-type lectin receptors that bind to *C. albicans* [40]. In addition, murine studies using DC-specific knockout of SHP-2 or small molecule inhibitors showed that SHP-2 is important for DC migration to the secondary lymph nodes, which is essential for the induction of CTL responses, as this is where DCs cross-present antigen to naïve CD8^+^ T cells [41]. On the other hand, SHP phosphatases also can inhibit this process, as experiments in mice with CD11c^+^ specific knockout for SHP-1 or with 20 μM NSC-87877 showed that SHP-1 negatively regulates the activation of antigen specific CTLs in vivo and in vitro [28]. Mechanistically, this was attributed to reduced endosomal acidification in the absence of SHP-1, which could preserve the antigen for cross-presentation [28].

Our phosphoproteomics identified only 12 phosphotyrosine sites that were significantly altered after pathogenic stimulation, of which six became more phosphorylated. These include the well-known phosphorylation sites for activation of MAPK signaling MAPK13 Y182 and MAPK9 Y185 and it might well be that SHP-1/2 inhibition further increases their phosphorylation, and that this affects cross-presentation somehow. However, the canonical STAT3 activation site Y708 also became more phosphorylated upon pathogen recognition, and this could be required for cross-presentation because murine experiments with STAT3 specific knockout in CD103^+^ cDC1 showed that STAT3 inhibits DC maturation and regulation of anti-tumor immunity [42], and it has been shown that STAT3 is a substrate of SHP-1 [43].

The other six tyrosines became more dephosphorylated, potentially by SHP-1, SHP-2 or other tyrosine phosphatases. Interestingly, the dephosphorylated tyrosines that we identified in our screen include proteins that are involved in T-cell activation. For example, Plastin-2 (LCP1) is well known to regulate T-cell migration and formation or maintenance of the immunological synapse by regulating F-actin polymerization and binding to the integrin LFA-1 [44]. It might well be that plastin-2 exerts similar functions in DCs, and its dephosphorylation might result in dysfunctional interactions with T cells. However, although Plastin-2 is well known to be regulated by serine phosphorylation of multiple residues in its N-terminal regulatory domain, the functional consequences of tyrosine 214 dephosphorylation (located in an actin-binding domain) are still unknown. Another protein that becomes dephosphorylated upon pathogen recognition is NFAT2C. This protein is also involved in T-cell activation, because knockout of the NFAT2C coding gene in a murine transplantation model showed decreased CTL activity and less DC migration [45]. Another possibility is that SHP-1 and/or SHP-2 reduce the efficiency of antigen cross-presentation through RasGAP, because our phosphoproteomics data show that Y299 of docking protein downstream of tyrosine kinase 2 (DOK2) also becomes dephosphorylated in LPS activated DCs. DOK2 is a tumor suppressor that plays well-known roles in cancer [46] and Y299 has been implicated in activation of RasGAP [47]. In vitro and in vivo experiments of viral infection showed that CD8+ T cells with DOK2 silenced had a slightly reduced magnitude of virus-specific effector CD8+ T cell expansion, but increased induction of cell surface TCR, granzyme B and tumor necrosis factor (TNF). However, DOK2-deficient effector CD8^+^ T cells showed a strongly reduced long-term survival, and this resulted in impaired generation of memory CD8^+^ T cells. These findings indicate that DOK2 negatively regulates the overactivation of CD8^+^ T-cells and promotes the formation of memory cells through RasGAP signaling [48].

In conclusion, our data indicate that SHP-1/2 inhibition blocks antigen cross-presentation of cancer antigens NY-ESO-1 and gp100 by human macrophages and DCs. This finding contrasts previous findings in mouse CD11c^+^ DCs [28], contributing to the notion that cross-presentation pathways vary among species, cell types and antigens [1]. Moreover, our findings contribute to the emerging understanding that SHP-1/2 does not only inhibit in the immune system, but also plays important stimulatory roles for both adaptive and innate immune responses [38-41]. These roles should be considered in therapies targeting SHP-1/2, especially in combination with immunotherapies.

## Data availability

The data that support the findings of this study are available from the corresponding author upon reasonable request.

