## Supplemental figure 1 for "Inhibition of SHP-1 /2 blocks antigen cross-presentation by human macrophages and dendritic cells"

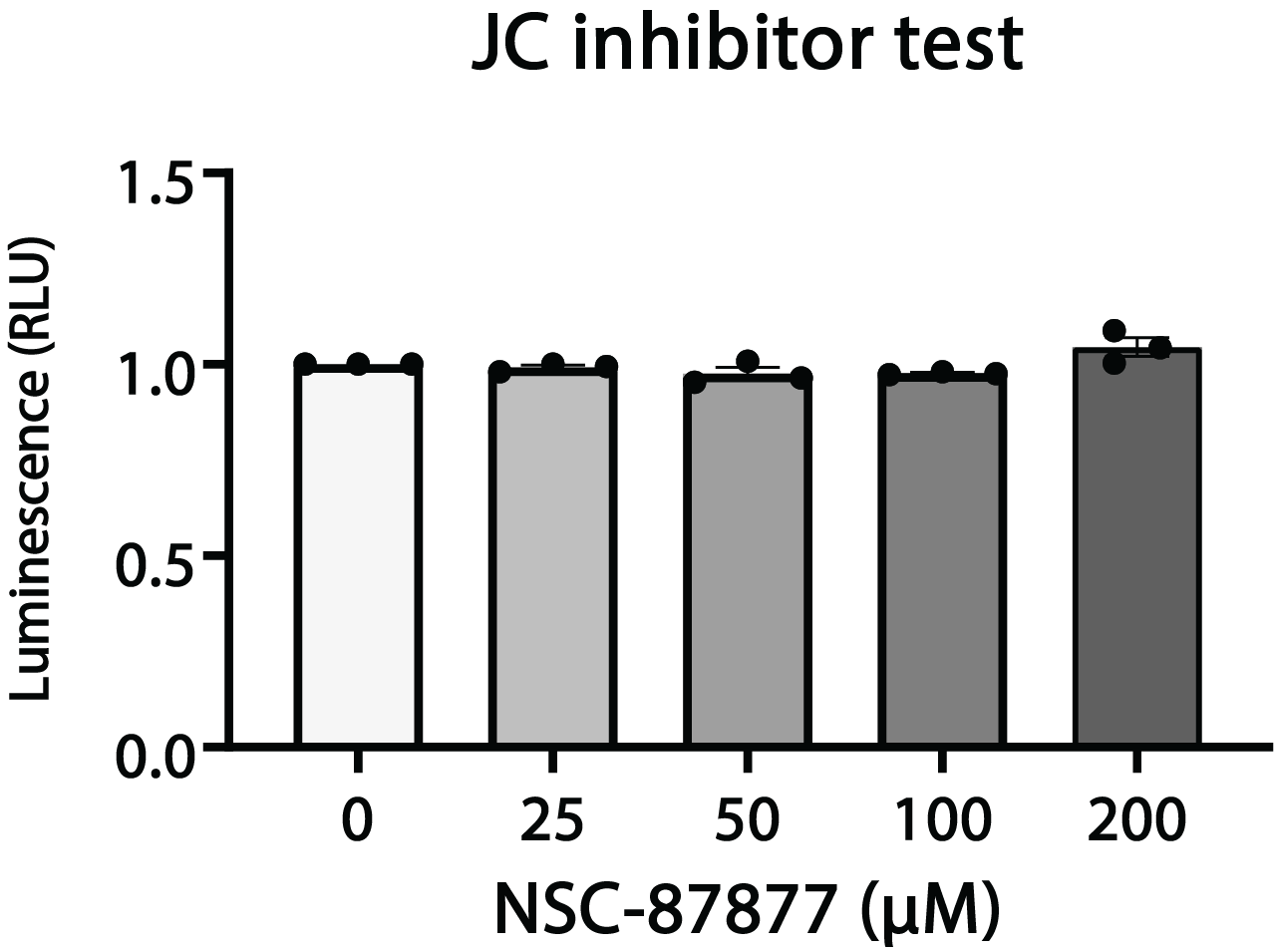

**Supplemental figure 1. Response of Jurkat reporter T-cell line is not affected by SHP1/2 inhibition.** The Jurkat T cells were treated with stimulated with PMA and ionomycin and different concentrations of SHP1/2 inhibitor NSC-87877 for 30 minutes to check the effect of the inhibitor on luciferase production.
