## Supplemental figure 2 for "Inhibition of SHP-1 /2 blocks antigen cross-presentation by human macrophages and dendritic cells"

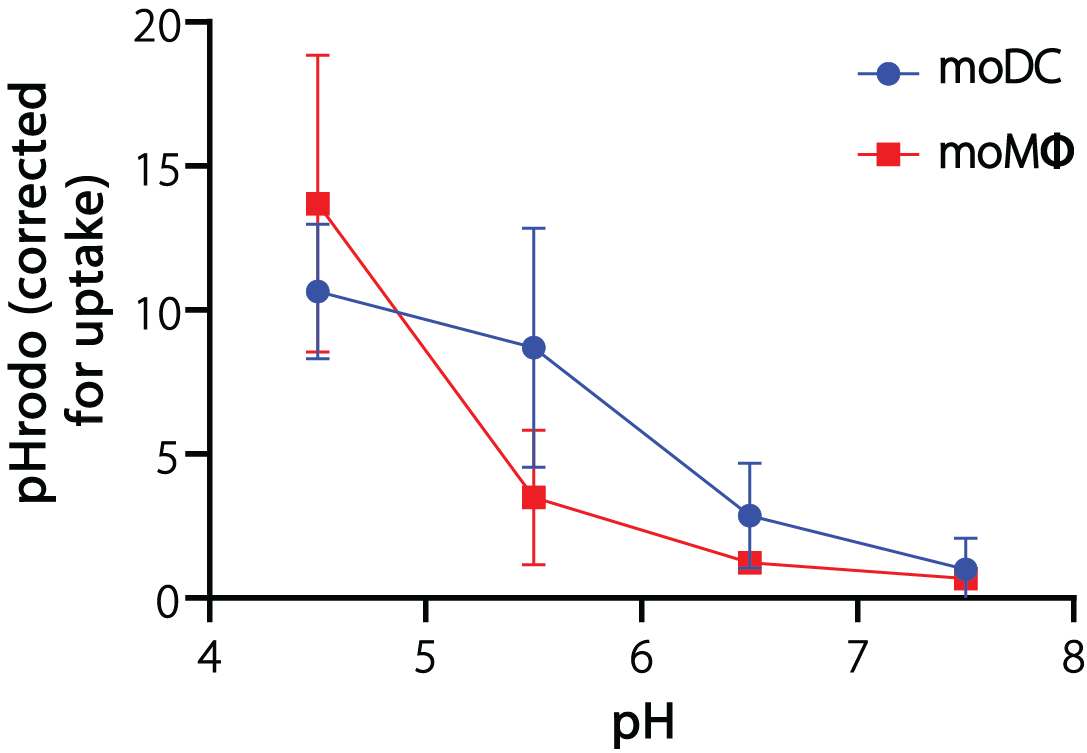

**Supplemental figure 2. pH calibration curve.** Monocyte-derived dendritic cells and macrophages were treated with pHrodo-labeled dextran for 30 min. To correct for uptake, cells were simultaneously treated with Alexa Fluor 647-labeled dextran. After treatment, cells were washed and incubated in buffers of different known pH and fluorescence intensity was determined by flow cytometry.
